# Gestational insulin resistance is mediated by the gut microbiome-indoleamine 2,3-dioxygenase axis

**DOI:** 10.1101/2021.07.21.453234

**Authors:** Medha Priyadarshini, Guadalupe Navarro, Derek J Reiman, Anukriti Sharma, Kai Xu, Kristen Lednovich, Christopher R Manzella, Md Wasim Khan, Barton Wicksteed, George E Chlipala, Barbara Sanzyal, Beatriz Penalver Bernabe, Pauline M Maki, Ravinder K Gill, Jack Gilbert, Yang Dai, Brian T Layden

## Abstract

**Background and aims:** Normal gestation involves reprogramming of maternal gut microbiome (GM) that may contribute to maternal metabolic changes by unclear mechanisms. This study aimed to understand the mechanistic underpinnings of GM – maternal metabolism interaction.

**Methods:** The GM and plasma metabolome of CD1, NIH-Swiss and C57BL/6J mice were analyzed using 16S rRNA sequencing and untargeted LC-MS throughout gestation and postpartum. Pharmacologic and genetic knockout mouse models were used to identify the role of indoleamine 2,3-dioxygenase (IDO1) in pregnancy-associated insulin resistance (IR). Involvement of gestational GM in the process was studied using fecal microbial transplants (FMT).

**Results:** Significant variation in gut microbial alpha diversity occurred throughout pregnancy. Enrichment in gut bacterial taxa was mouse strain and pregnancy time-point specific, with species enriched at gestation day 15/19 (G15/19), a point of heightened IR, distinct from those enriched pre- or post- pregnancy. Untargeted and targeted metabolomics revealed elevated plasma kynurenine at G15/19 in all three mouse strains. IDO1, the rate limiting enzyme for kynurenine production, had increased intestinal expression at G15, which was associated with mild systemic and gut inflammation. Pharmacologic and genetic inhibition of IDO1 inhibited kynurenine levels and reversed pregnancy-associated IR. FMT revealed that IDO1 induction and local kynurenine levels effects on IR derive from the GM in both mouse and human pregnancy.

**Conclusions:** GM changes accompanying pregnancy shift IDO1-dependent tryptophan metabolism toward kynurenine production, intestinal inflammation and gestational IR, a phenotype reversed by genetic deletion or inhibition of IDO1.

## Introduction

Dynamic physiological and metabolic changes occur throughout pregnancy ^1^. Aberrations in these processes can result in pregnancies complicated by hyperglycemia and/or gestational diabetes mellitus, a harbinger of future diabetes in the mother and metabolic disease in the offspring. In early gestation, the maternal body is in an anabolic state with increased insulin sensitivity promoting adipose lipid stores. In contrast, late gestation is a catabolic state with reduced insulin sensitivity resulting in increased circulating free fatty acids and decreased fasting glucose levels ^2^. These changes are considered beneficial for growth of the fetus and preparing the maternal body for lactation ^3^.

Insulin resistance (IR) of normal pregnancies is multifactorial, with poorly understood underlying mechanisms ^1^. Recently, the composition of the gut microbiome (GM) was shown to be altered during pregnancy ^4^ and associated with altered metabolic features in the mothers ^5^. Koren et al. ^5^ showed that the third trimester maternal GM resembles the dysbiotic microbiome of the metabolic syndrome and promotes metabolic syndrome-like features in pseudo germ-free (GF, antibiotic treated) mice. Whether these changes are pre-requisite to the physiological adaptations during pregnancy is unclear. Moreover, there is lack of consensus on the changes in the microbiota of mothers ^5–7^, and how ethnicity, geographical location, dietary habits and environmental factors can affect the GM ^8^.

Thus, highly controlled mouse studies can clarify the exact nature of GM changes during pregnancy and their influence on metabolism during pregnancy. Significant strain and vendor differences in the GM populations of mice ^9, 10^ were leveraged in this study to capture changes in gut microbial and metabolomic features through pregnancy. This allowed identification of a metabolite, kynurenine, to be increased during pregnancy and demonstration of its importance in pregnancy-associated metabolic physiology.

## Results

### Mouse genetic background influences the metabolic response to pregnancy

Physiological and metabolic responses of the three mouse strains (C57Bl6/J, CD1, and NIH-Swiss) were similar and as expected ^1^, with early increases in adiposity and circulating glucose, followed by IR and a concomitant reduction in adipose stores to fuel fetal growth and maternal lactation (Fig. S1). These data illustrate that pregnancy induces specific physiological and metabolic responses with strain-specific variations.

### Strain specific gut microbial composition changes and phenotypic associations during pregnancy

To systematically explore how pregnancy influences the GM, we investigated the fecal microbial communities of the three strains of mice where each strain is expected to have a unique highly conserved gut flora ecosystem ^10^. Alpha diversity, calculated using Shannon index (accounts for richness and evenness), was significantly different between pregnancy stages and between mouse strains (*P_FDR_*<.05, non-parametric, Fig. 1a). Each strain had a unique pattern through pregnancy with NIH-Swiss mice more microbially diverse than CD1 or C57BL6 mice (*P_FDR_*<.05) (Fig. 1a). Microbiota differences between mouse strains and across pregnancy stage were calculated by beta diversity using the unweighted (Fig. S2a) and weighted UniFrac distance metrics (Fig. 1b). Arrayed using non-metric multidimensional scaling (NMDS) plots, these data demonstrated significantly differential clustering pattern by mouse strain (*P_FDR_*<.05) (Fig. 1b), but not significant clustering by pregnancy time-point (Fig. 1c, S2a). As, unweighted UniFrac distance metrics, unlike weighted UniFrac, equally weighs both rare and dominant species, these results suggest that the three mouse strains do not exhibit commonalities in low-relative abundance taxa.

**Figure 1.**
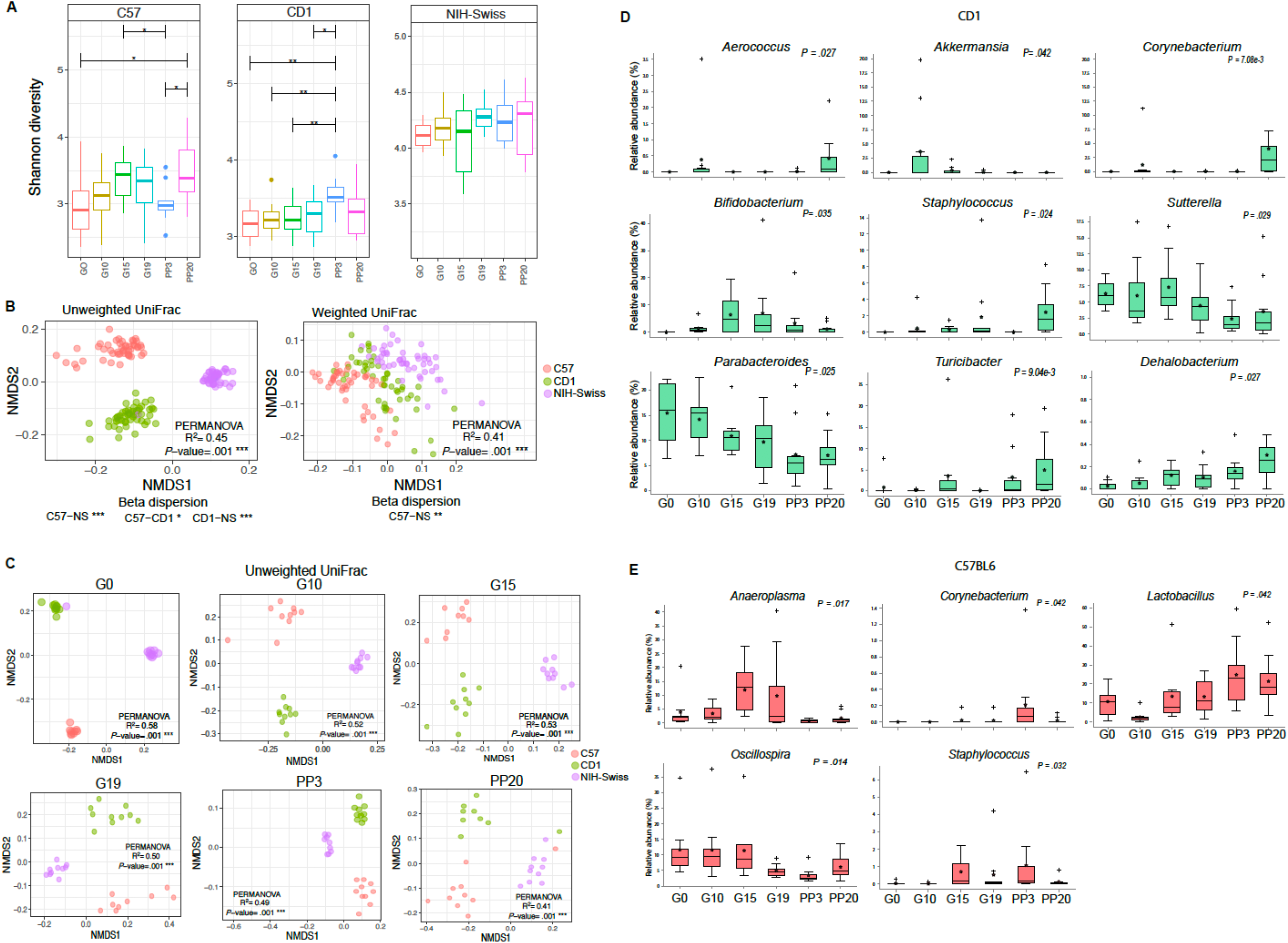
Pregnancy impacts fecal bacterial diversity and select bacterial taxa. **(a)** Box plots of alpha diversity (Shannon index); **(b-c)** non-metric multidimensional scaling plots (NMDS) based on weighted and unweighted UniFrac dissimilarity matrix between mouse strains **(b)** and across pregnancy stages **(c)**; **(d-e)** multi-group non-parametric analyses of composition of microbiome (ANCOM) followed by Mann-Whitney U test indicating differentially abundant genera at various stages of pregnancy in CD1 **(d)** and C57BL6/J **(e)** mice. *p-values* (Benjamini-Hochberg FDR correction) by Kruskal Wallis **(a)**, permutational multivariate analysis of variance (PERMANOVA) **(b-c)**.

To identify the compositional variations at both genus and Exact Amplicon Sequence Variant (ESV) level between pregnancy phases in each strain, we performed multi-group ANCOM analysis (Fig. 1 d-e; S2b). In CD1 mice *Aerococcus, Akkermansia, Bifidobacterium, Sutterella, Turicibacter*, and *Dehalobacterium* were significantly enriched in pregnancy, *Parabacteroides* was most abundant at gestation day (G) 0 and reduced at G15 to postpartum day (PP) 20, and *Corynebacterium* and *Staphylococcus* were enriched at PP20 (for all *P_FDR_*<.05; Fig. 1d). In C57/BL6, *Lactobacillus* increased significantly over the gestational period while *Aneroplasma* was significantly more abundant at G15 and G19 compared to other time-points (for both *P_FDR_*<.05; Fig. 1e). In NIH-Swiss, *Candidatus arthromitus* peaked at G15 (*P_FDR_*<.05) (Fig. S2b). These ESVs when arranged in modules based on co-occurrence exhibited distinct correlations with above parameters (**supplemental information, SI**; Fig. S2c).

Using K clustering (Fig. S2d**, SI**), mice were grouped based on the gestational time, with G15/G19 as one group and G0, G10, PP3 and PP20 as the other. Meta-Signer, a tool that produces a robust ranked list of the taxa based on multiple machine learning methods, was used to identify the most discriminative taxa between the two groups ^11^. The top 10 taxa on the ranked list could significantly separate the G15/19 and non-G15/19 groups (*P=*.006) (Fig. 2a), where *Anaeroplasma*, *Candidatus Arthromitus, Sutterella,* and family Lachnospiraceae showed higher average abundance in G15/G19 mice; and *Bacteroides, Corynebacterium*, *Acinetobacter*, and *Lactococcus* showed a lower average abundance (Fig. 2b). Additionally, these taxa were found with high level of co-occurrence based on Spearman’s rank correlation coefficients (Fig. 2c, *P*<.05) that was consistent across mouse strains. Interestingly, the full microbe set could also significantly separate the G15/19 and non-G15/19 groups (*P*=.009) (Fig. S2e). These data corroborate with earlier reports on distinctive shift in GM in later pregnancy in both humans and mice ^12^.

**Figure 2.**
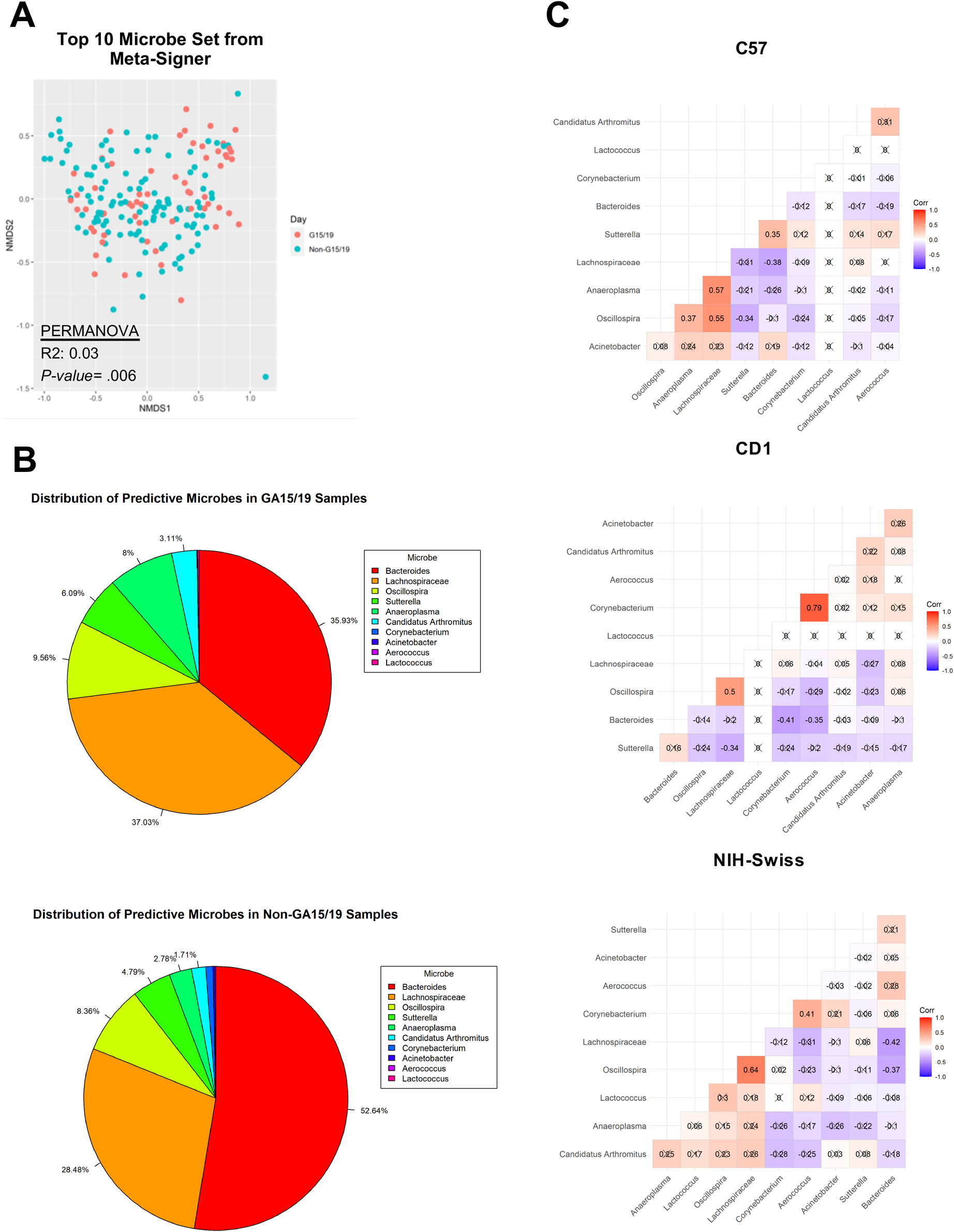
Microbial taxa predict gestational age. **(a)** NMDS plot using the top 10 microbes identified by Meta-Signer across all strains; **(b)** distributions of the top 10 microbes within G15/19 and non-G15/19 samples; **(c)** Spearman correlation values between the top 10 microbes’ abundance values within strains.

### Plasma metabolome is unique to mouse gestational age and strain

Metabolomic analyses by untargeted LC-MS revealed substantial changes in maternal metabolism in pregnancy, especially in later stages. Overall, there were 39,323 and 2,636 features in the positive and negative mode datasets, respectively, after filtering to remove features that were present in less than 40% of the samples. First, PERMANOVA analysis identified >700 significantly changed metabolites in NIH-Swiss and CD1 mice (*P*<.05) and 974 such metabolites in C57Bl6/J mice (*P*<.05) during the course of gestation and postpartum (Fig. S3a). Further, even though the metabolome showed strong differences in strains at each time-point (Fig. S3b), it was ordered similarly by the gestational state within each strain (Fig. 3a). Using MetaboAnalyst ^13^ and Mummichog algorithm ^14^, we next identified top pathways significantly altered in G15/19 phase (**Tables S1-3**). As expected, several pathways of fatty acid metabolism, amino acid metabolism, steroid hormone biosynthesis and vitamin metabolism were enriched in the G15/19 phase. Among the amino acid metabolism pathways, tryptophan metabolism was significantly affected in all three strains (Fig. 3b, S3c&f).

**Figure 3.**
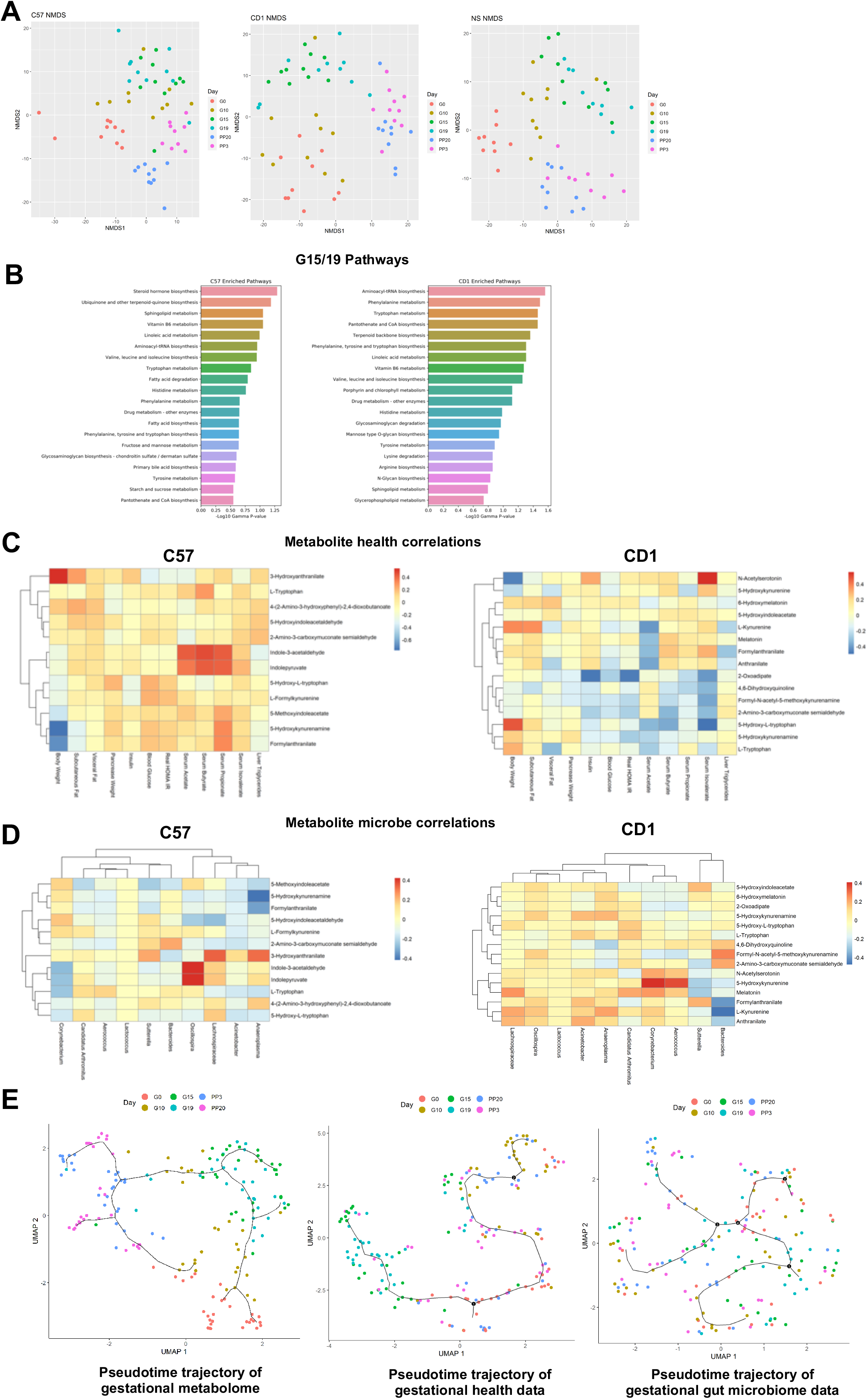
Plasma metabolome strongly clusters by gestational age and mouse strain. **(a)** NMDS plots of metabolomic data within each strain colored by gestational state; **(b)** the top 20 most significantly enriched pathways based on the mummichog analyses on the differential metabolites for C57BL/6J (left) and CD1 (right); **(c)** Spearman correlation analyses between the top 10 most significant metabolites from the tryptophan metabolism pathway that were able to be mapped using mummichog candidates and both the health parameters and **(d)** the top 10 microbes identified from Meta-Signer for the C57 and CD1 strains; **(e)** gestational trajectories identified using Monocle3 using UMAP reduction and Louvain clustering.

As the GM and plasma metabolome could order the gestational time-points (Fig. 2a; S2e; 3a; S3b), we next determined the association between the metabolomic, gut microbial and health profiles. We observed both common and strain specific associations in plasma metabolites with pregnancy metabolic parameters (Fig. 3c; S3d). Tryptophan metabolites like L-kynurenine and its derivatives (e.g. 3-hydroxyanthranilate) correlated with body weight, adipose and HOMA-IR as observed in other human metabolic states ^15^. We also found previously unknown associations of 5-hydroxykynurenamine, a serotonin antagonist ^16^, and the melatonin precursor, N-acetylserotonin with pregnancy body weight. With well appreciated roles of melatonin and peripheral serotonin in energy metabolism and pregnancy ^17, 18^, these associations suggest additional points of regulation of pregnancy adaptations. Associations were also observed between the top 10 G15/19 predictive microbes and metabolites (Fig. 3d, S3e), including between tryptophan metabolites and microbial features associated with obesity, diabetes and inflammation e.g. Lachnospiraceae and *Sutterella*.

To measure potential shifts of the metabolic profiles, microbiome, and plasma metabolome during the course of gestation and postpartum, Monocle 3 (originally developed for single cell RNA-Seq data to construct cell trajectories across pseudotime) ^19^ was used to visualize potential trajectories based on their profiles over pseudotime (Fig. 3e). The G15/G19 mice were well clustered based on the metabolic parameters and metabolomic data, with the latter giving an even tighter cluster. Furthermore, in the pseudotime trajectory of gestational metabolome data, mice were grouped such that the gestational stages can be traced in temporal order, G0/G10, G15/G19 and PP3/PP20, a similar observation that has been recently described in human pregnancy ^20^. The microbiome data, on the other hand, provided a mixed clustering pattern regarding the gestational stages.

### The tryptophan metabolite, kynurenine, is elevated during pregnancy

Our untargeted metabolomics data revealed enrichment of the tryptophan catabolic pathway in pregnancy in all three strains of mice. Since tryptophan metabolism is strongly associated with GM, and to understand the potential metabolic role of pregnancy related tryptophan metabolism, we employed targeted metabolomics to assess this pathway (Fig. 4a). Plasma kynurenine was elevated in each mouse strain during the IR phase of pregnancy and returned to pre-gestation levels postpartum (Fig. 4b). Serotonin decreased during pregnancy in each mouse line, while tryptophan only decreased in CD1 mice (Fig. 4b) and indoleacetic acid remained unchanged (Fig. S4a). With known roles of serotonin in pregnancy induced adaptation of beta cells ^18^, we explored the levels of these metabolites in the pancreas, observing tryptophan, kynurenine and serotonin to be significantly increased in the pancreas during pregnancy although tryptophan levels remained unaltered in NIH-Swiss mice (Fig. S4b).

**Figure 4.**
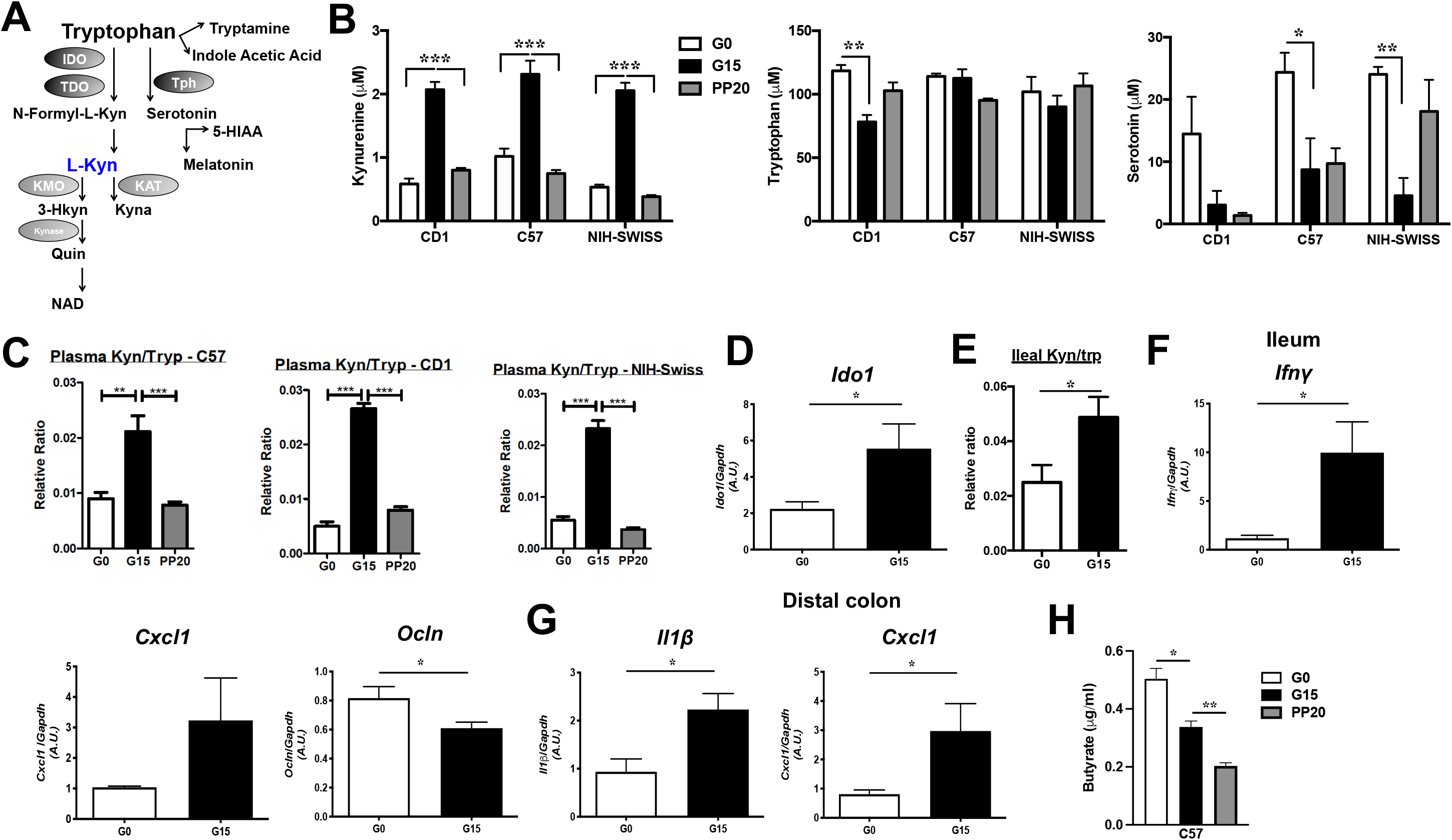
The tryptophan metabolite, kynurenine, is elevated during pregnancy. **(a)** Schematic of the tryptophan metabolism pathways; **(b)** plasma kynurenine, tryptophan and serotonin levels and **(c)** plasma kynurenine/tryptophan ratio (kyn/tryp) in C57BL/6J, CD1 and NIH-Swiss mice at G0, G15 and PP20; **(d-f)** in the ileum of C57BL/6J mice at G0 and G15, *Ido1* mRNA expression **(d)**, kynurenine/tryptophan ratio **(e)** and mRNA expression of *Ifnγ*, *Cxcl1*, and *Ocln* **(f)** and **(g)** mRNA expression of *Il1β* and *Cxcl1* in DC of same mice; **(h)** plasma butyrate levels in C57BL/6J, CD1 and NIH-Swiss mice at G0, G15 and PP20 (n=10 per time-point). Data are mean±SE, n=5-7 per time-point analyzed by one-way ANOVA with Tukey’s post-hoc analysis (**b-c)**, two-tailed Student’s unpaired *t* test **(d-g)**, one-way ANOVA (Mixed effects analysis) (**h**). \**P*<.05, \*\**P*<.01, \*\*\**P*<.0005, \*\*\*\**P*<.0001. IDO-Indoleamine 2,3-dioxygenase; TDO-Tryptophan 2,3-dioxygenase; Tph-Tryptophan hydroxylase; KAT-Kynurenine aminotransferase; KMO-Kynurenine 3-Monooxygenase; Kynase-Kynureninase; N-Formyl-L-Kyn-N-Formyl-L-Kynurenine; L-Kyn-Kynurenine; HIAA-5-Hydroxyindoleacetic acid; Quin-Quinolinic acid; Kyna-Kyneurenic acid; 3-Hkyn-3-Hydroxykynurenine.

Elevated kynurenine suggested increased tryptophan catabolism and flux through the kynurenine pathway; therefore, we computed kynurenine to tryptophan ratio (K/T) in the plasma and the pancreas. K/T was significantly higher during pregnancy in plasma and pancreas in each mouse strain (Fig. 4c, Fig. S4d). K/T is reflective of indoleamine-2,3-dioxygenase 1 (IDO1) activity, the rate limiting enzyme of tryptophan degradation in kynurenine pathway ^21^ which is known to be expressed in immune, adipose, and intestinal cells ^22^. During pregnancy (G15), *Ido1* mRNA expression increased in the ileum of C57BL/6 mice (Fig. 4d), without a difference in the colon, and trended higher in the ileum and was significantly higher in the colon of CD1 mice (Fig. S4e). Consequent to this increased *Ido1* expression, K/T was significantly higher in C57BL/6 ileal tissue at G15 (Fig. 4e). Subsequent experiments used C57BL/6 mice for simplicity and the availability of a IDO1 knockout model on the C57Bl/6 background.

### Low grade inflammation in pregnancy modulates gut IDO1 activity

Normal pregnancy is a state of low-grade inflammation, specifically at gut mucosal surfaces and tissues like adipose with a build-up of proinflammatory cytokines ^5, 23^ and inflammation upregulates IDO1 expression in the intestinal epithelium ^24^. Thus, we assessed mRNA expression of cytokine genes in G0 and G15 ileum and distal colon (DC) mucosa. At G15, ileal expression of *Ifnγ*, the main inducer of IDO1 ^22^, was significantly high while expression of the pro-inflammatory chemokine, *Cxcl1*, trended high (Fig. 4f). Expression of pro-inflammatory *Il1β* and *Cxcl1* increased significantly in DC (Fig. 4g). The tight junction (TJ) proteins of the intestinal epithelium, along with other protein networks, guard the entry across the epithelium, and their expression is negatively impacted by gut inflammation ^25^. Accordingly, expression of occludin (*Ocln*), a TJ protein, reduced significantly in G15 ileum (Fig. 4f). Additionally, short chain fatty acids (SCFAs) modulate gut IDO1 expression, where butyrate can downregulate IDO1 expression in intestinal epithelial cells ^26^. Of note, plasma levels of total SCFAs and butyrate declined significantly from G0 to G15 in C57BL/6 mice (Fig. S4f; Fig. 4h).

### IDO1 activity affects pregnancy IR

As kynurenine was specifically elevated at the height of gestational IR, we next assessed the role of kynurenine/IDO1 in pregnancy first by pharmacologically inhibiting IDO1 utilizing specific inhibitor L-1Methyltryptophan (L1MT), by supplementing the drinking water ^27^. L1MT treated mice did show reduced plasma and ileal tissue K/T (Fig. 5a & 5b), particularly during pregnancy. No differences were observed in body weight gain during pregnancy and number of pups in control and L1MT treated mice (data not shown). Surprisingly, L1MT treated mice displayed glucose intolerance prior to pregnancy (Fig. 5c). Though not noted before ^27^, it may reflect indirect effects of L1MT or metabolic effects of sucralose from Splenda ^28^. Contrary to this, at G15, L1MT treated mice exhibited improved glucose tolerance (Fig. 5c) along with reduced IR (Fig. 5d) and corresponding increase in insulin signaling (phosphorylated-Akt) in the soleus muscle (Fig. 5e, S5a).

**Figure 5.**
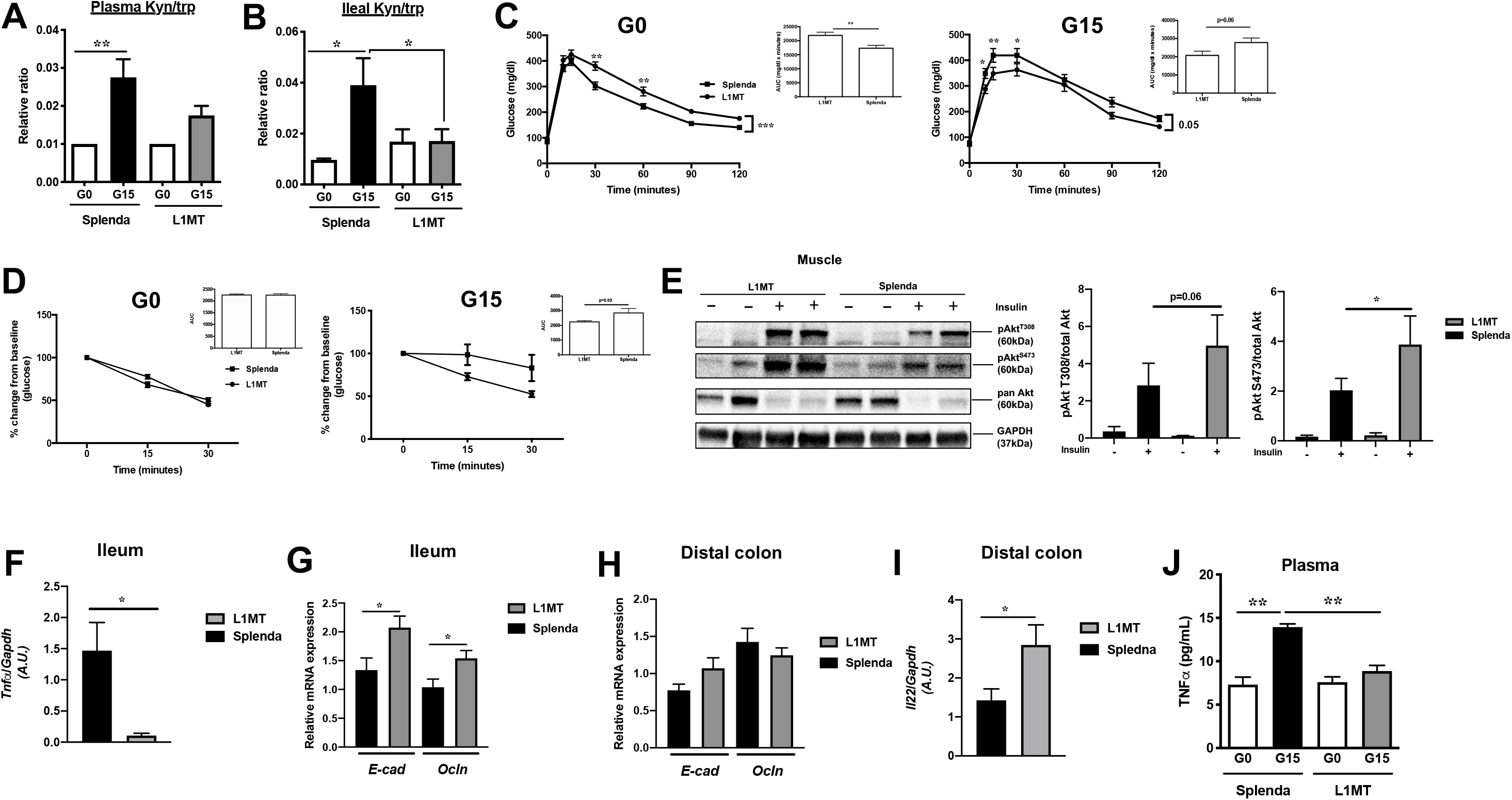
IDO1 activity affects pregnancy induced IR. 1-methyl-L-tryptophan (L1MT) treated and control mice were analyzed for kynurenine/tryptophan ratio (kyn/trp) **(a)** in the plasma, and **(b)** in the ileum; **(c-d)** intraperitoneal glucose and insulin tolerance tests, respectively, at G0 (*left, n=4-5*) and G15 (*right, n=5-7*) (inset, area under curve, AUC); **(e)** insulin signaling (p-Akt-T308 and p-Akt-S473) in soleus muscle with corresponding densitometric analysis (n=4-5); mRNA expression of **(f)** *Tnfα*, **(g)** *E-cad*, *Ocln* in the ileum and **(h)** *E-cad*, *Ocln*, **(i)** *Il22* in the DC, at G15 (n=5-7); **(j)** plasma Tnfα levels at G0 (n=3) and G15 (n=5-7). Data are mean±SE and analyzed by one-way ANOVA with Tukey’s post-hoc analysis **(a-b)**, two-tailed Student’s unpaired *t* test (insets **c-d**, **e-h)**, two-way ANOVA with Tukey’s post-hoc analysis (**c, d**). \**P*<.05, \*\**P*<.01, \*\*\*\**P*<.0001.

IDO1 activity inhibition partially protected against pregnancy induced gut inflammation, in the ileum, noted by significantly lower expression of pro-inflammatory *Tnfα* ^29^, modest but significant upregulation of barrier proteins, *Ocln* and E-cadherin (*E-cad*) and modest increase in expression of claudin 4 (*Cldn 4*) in the ileum of L1MT treated mice (Fig. 5f-g, S5b). The expression of other tested proinflammatory transcripts showed modest changes (*Il1β* and/or *Il6*) in L1MT treated mice both in ileum and DC (Fig. S5b). In DC, no differences were observed in the expression of proinflammatory cytokines or gut barrier proteins (Fig. 5h; Fig. S5b), although, expression of the anti-inflammatory cytokine, *Il22*, was significantly upregulated in L1MT treated mice (Fig. 5i). Contrasting controls, plasma levels of Tnfα, known to be increased in insulin resistant phase of pregnancy and obesity ^1^ were significantly reduced in L1MT treated mice (Fig. 5j).

Thus, IDO1 inhibition reduced the systemic and locally increased K/T, moderately enhanced gut barrier and reduced IR. However, due to dysregulated glucose tolerance prior to pregnancy (Fig. 5c) and the unknown extent and sustainability of pharmacological IDO1 inhibition with L1MT, we sought to confirm these findings using a more definitive model, mice with genetic knockout of IDO1.

### Genetic IDO1 deficiency protects against pregnancy induced IR

Mice lacking IDO1 throughout the body (IDO-KO) were characterized relative to control, C57Bl/6J mice, prior to gestation and at G15. Contrasting controls, in IDO-KO mice, K/T ratios was markedly suppressed in plasma at G15 and in ileum at G0 and G15 (Fig. 6a-b). Compared to controls, IDO-KO pregnant mice displayed significant protection against glucose intolerance (Fig. 6c-d) and pregnancy induced IR (Fig. 6e) with correspondingly higher insulin signaling (phosphorylated Akt) in adipose (Fig. 6f; Fig. S5c).

**Figure 6.**
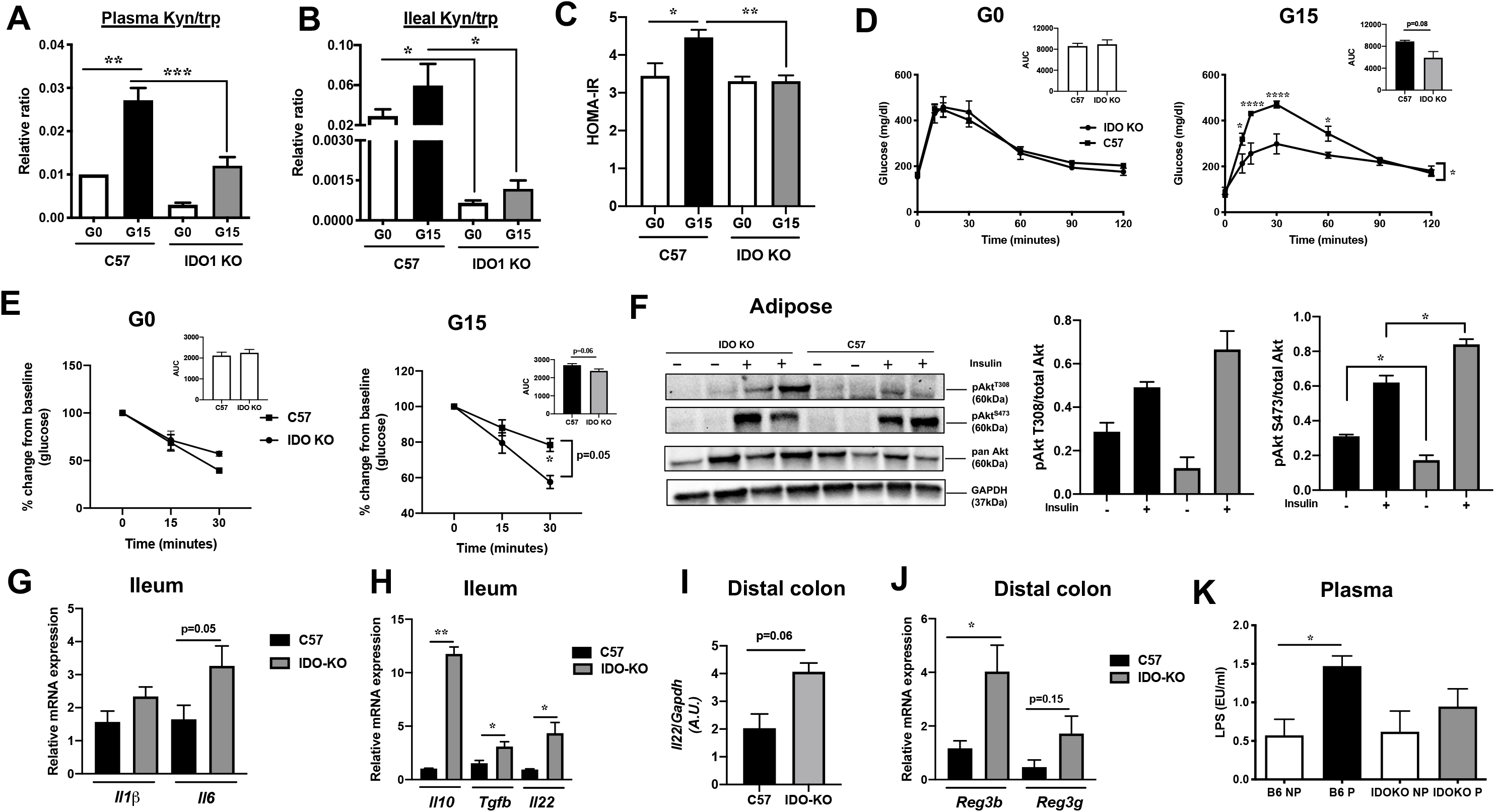
Genetic deficiency of IDO1 protects against pregnancy induced IR. IDO1 KO and control mice were analyzed for kynurenine/tryptophan ratio (kyn/trp) in plasma **(a)** and in ileum **(b)**; **(c)** HOMA-IR, **(d-e)** intraperitoneal glucose and insulin tolerance tests, respectively, at G0 (*left, n=4-5*) and G15 (*right, n=5-7*) (inset, AUC); **(f)** insulin signaling (p-Akt-T308 and p-Akt-S473) in subcutaneous adipose with corresponding densitometric analysis (n=4-5); mRNA expression of **(g)** *Il1β*, *Il6* and **(h)** *Il10, Tgfβ*, *Il22* in the ileum and of **(i)** *Il22* and **(j)** *Reg3b and Reg3g* in the DC at G15 (n=5); **(k)** plasma LPS levels at G0 (n=3) and G15 (n=5). Data are mean±SE analyzed by one-way ANOVA with Tukey’s post-hoc analysis **(a-b)**, two-tailed Student’s unpaired *t* test (insets **d**-**e & f-j)**, and two-way ANOVA with Tukey’s post-hoc analysis (**c-e, k**). \**P*<.05, \*\**P*<.01.

As well known, IDO1 has immunoregulatory effects ^22^. No differences, however, were noted in expression of proinflammatory *Cxcl1*, *Ifnγ*, or *Tnfα* transcripts in ileum or DC (data not shown) except increased *Il6* in the ileum of IDO-KO mice (Fig. 6g). As the activity of IDO1 sustains an immunostimulatory state by inhibiting IL-10 production ^30^, we analyzed the expression of anti-inflammatory cytokine, *Il10*, observing in the ileum of IDO-KO mice 11-fold enhanced expression of *Il10* along with upregulation of another anti-inflammatory cytokine, *Tgfβ* ^31^ (Fig. 6h). Additionally, expression of *Il22*, a cytokine promoting gut homeostasis and tissue regeneration ^32^ was increased in both ileum and DC of IDO-KO mice (Fig. 6h-i). We further found increase of Il22 target genes such as antimicrobial proteins, regenerating islet-derived protein (*Reg*) *3g* and *Reg3b* in DC of IDO-KO mice (Fig. 6j). Consequent to the anti-inflammatory milieu in IDO-KO gut, plasma lipopolysaccharide (LPS) levels were not elevated at G15 relative to G0, in contrast to control mice (Fig. 6k).

### Gut microbiome-IDO1 axis mediates pregnancy IR

GM can influence kynurenine pathway either by modulating inflammation which impacts IDO1 expression or by affecting tryptophan availability ^22^. To test the involvement of GM in IDO1 mediation of pregnancy IR, we explored whether fecal microbial transfer (FMT) from pre-gestation and G15 C57Bl/6J mice to antibiotic induced pseudo-germ free (GF) C57Bl/6J mice could recapitulate the respective G0 and G15 pregnancy phenotypes (Fig. 7a) where G15 time point, unlike G0, is characterized by increased IR, K/T ratio, gut IDO1 expression and inflammation. While no weight difference emerged (Fig. 7b), G15 recipients, compared to G0 recipients, exhibited a trend toward higher IR, increased ileal K/T and mRNA expression of *Ido1*; and the expression of inflammation marker lipocalin 2 (*Lcn2*) was significantly higher (Fig. 7c-f). These data suggest that G15 GM may be promoting tryptophan metabolism through IDO1 and the IR phenotype.

**Figure 7.**
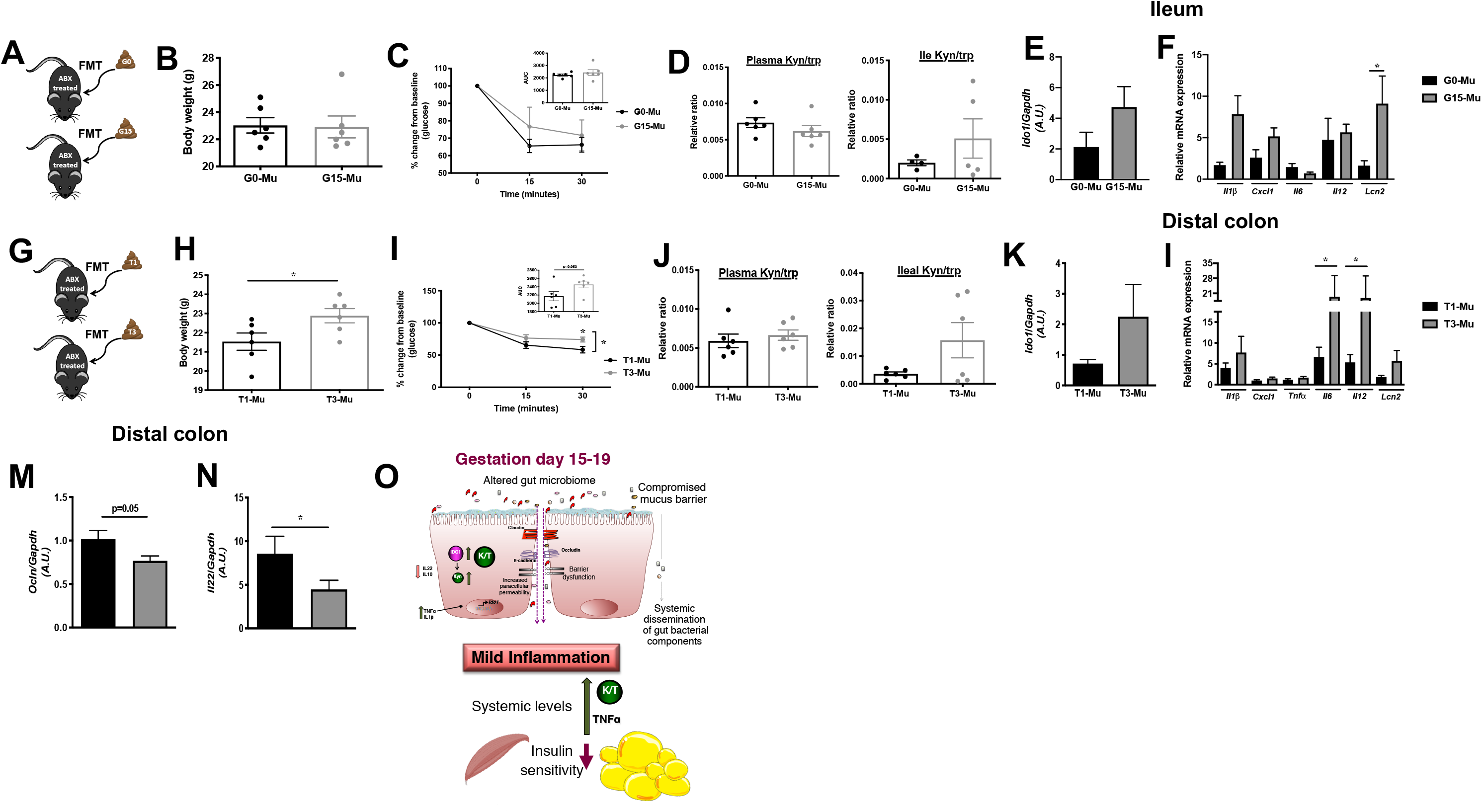
Transfer of pregnancy associated gut microbiota impacts IR, IDO1 levels, and intestinal inflammation. Schematic of fecal microbial transfer (FMT) to antibiotic (Abx) treated mice **(a, g)**; body weight **(b, h)**; insulin tolerance test (inset, AUC) **(c, i)**; kynurenine/tryptophan ratio (kyn/trp) in plasma (*left*) and ileum (*right*) **(d, j)**; mRNA expression of *ldo1* **(e, k)**, of proinflammatory cytokines and anti-microbial peptide *Lcn2* **(f, l)** in the ileum of G0 and G15 and in DC of T1 and T3 recipients, respectively; mRNA expression of **(m)** *Ocln*, **(n)** *Il22* in DC of T1 and T3 recipients. Shown in **(o)** is a summary: gut microbial change and mild inflammatory milieu at gut mucosal surfaces drives increase in gut IDO1 expression/activity (kynurenine/tryptophan ratio (K/T)) and shifts tryptophan metabolism to enhanced kynurenine (kyn) production in insulin resistant phase of pregnancy. n=6 per group, data are mean±SE analyzed by two-tailed Student’s unpaired *t* test (insets **b, d**-**f, h, j-n)** and two-way ANOVA with Tukey’s post-hoc analysis (**c, i**). \**P*<.05.

To identify if a similar mechanism in human pregnancy occurs, we transferred first trimester (T1) and third trimester (T3) fecal microbiota samples to pseudo-GF mice (Fig. 7g). T3 recipients displayed significantly increased body weight and IR and trend toward increased ileal K/T (Fig. 7h-j). Analysis of DC of T3 recipients revealed a trend toward higher *Ido1* mRNA expression (Fig. 7k), and significant differences in gut inflammation evident from significantly elevated *Il6* and *Il12*, higher *Lcn2*, and declined *Ocln* and *Il22* transcripts (Fig. 7l-n). As apparent, the transfer of the phenotype by FMT generates some expected features, but not all, such as increased in plasma K/T, which may be due to insufficient/partial colonization by GM.

## Discussion

The GM serves as crucial auxiliary in nutrition acquisition, maintaining gut homeostasis and immune programming playing important role in host health and disease ^33^. During pregnancy, GM impacts maternal and fetal health ^5, 34^. However, GM modulation of maternal adaptations to pregnancy are poorly understood. Through our comprehensive approach, we have discovered a role for kynurenine in gestational metabolism. Our data describes: (1) GM changes, throughout the course of gestation and postpartum across three genetically unique mouse lines, possibly impacting gut inflammation; (2) these changes impact tryptophan metabolizing enzyme, IDO1 expression and possibly activity as indicated by higher kynurenine to tryptophan levels; and (3) the mouse and human gestational GM mediates transferable aspects of these metabolic changes. Through these changes, kynurenine levels are impacted which in turn impacts gestational metabolism, most notably insulin resistance.

Earlier studies on gestational GM changes in rodents have explored fewer pregnancy time-points ^35, 36^ and have additional GM modifying components, such as high fat diet, in the study design, making extrapolation of the results to normal pregnancy less clear ^35, 37^.

Choice of 3 different strains of mice (from different vendors) each with distinct pre-pregnancy microbiomes ^9, 10^, phenotypic characteristics and propensities for metabolic disorders ^38, 39^, have allowed us to uniquely capture genotypic and phenotypic diversity of the microbial population cohorts ^39^. Our GM profiling results recapitulated aspects of previously reported pregnancy-specific GM features during the course of gestation and post-partum including changes in alpha diversity ^5^ and relatively stable beta diversity ^40^, with distinctive phyla, and genus level changes at each gestational time-point ^5^. With progression of pregnancy, a drift towards GM features associated with obesity, IR, gestational diabetes mellitus (GDM) and a proinflammatory states occurs ^5, 23^. G15/19 stages accompanied similar changes, including specifically a rise in the family Lachnospiraceae ^27, 39, 41^ and the genera: *Sutterella* from phylum Proteobacteria ^42^, *Staphylococcus* from phylum Firmicutes ^7^, *Aneroplasma* from phylum *Tenericutes* ^43^ and a decline in *Bacteriodes* ^41^. Corroborating earlier findings, transfer of G15 or T3 gut microbes to pseudo-GF mice elicited aspects of gut inflammation and a mild IR ^5^. The gut microbial changes seen during pregnancy, thus, seem to impact maternal metabolic adaptation to pregnancy ^7, 34, 44^.

The plasma metabolome was extensively remodeled by pregnancy. While expected enrichment of pregnancy signature features at G15/19, additional gut microbial signature metabolites were modulated, for example SCFAs, phenylalanine metabolites and several tryptophan metabolites. These data indicated associations between plasma metabolome, pregnancy and GM. We explored one such relationship, of the tryptophan metabolite kynurenine. A small percentage of tryptophan is metabolized directly by the GM for production of indole derivatives ^45^. Alternatively, as our data suggest, GM can indirectly impact the major arm of tryptophan metabolism catalyzed by IDO1 where gut dysbiosis affects gut IDO1 levels and/or activity, thereby influencing kynurenine levels. Current understanding of the contribution of different gut bacteria to plasma metabolites is incomplete. Studies assessing the effect of selective colonization of pregnant mice with defined bacterial populations on plasma features of tryptophan catabolic pathway and other significantly changing metabolites will help deconvolute complex interaction between maternal and gut bacterial metabolism. Nonetheless, our data reveals a crosstalk between gut environment and maternal systemic circulation that uniquely impacts maternal metabolism.

While IDO1 is a well-known immunoregulatory enzyme ^22^, its activity has only recently been implicated in metabolic disorders like obesity and diabetes ^22, 27^. The data presented here suggests a role of IDO1 in pregnancy IR (Fig. 7o). In the presence of the pregnancy associated proinflammatory milieu, systemically and at gut mucosal surfaces (indicated by presence of dysbiotic gut bacteria), there is an increase in IDO1 expression and activity that drives the formation of kynurenine, a molecule with multiple effects on peripheral tissues ^27, 46, 47^ including the development of IR. IDO1 deficiency, in contrast, reduces kynurenine production promoting an anti-inflammatory gut environment. Lastly, the role of IDO1 in mediating pregnancy IR is associated, at least partially, with GM in both mice and human pregnancy.

Future steps include the delineation of specific features of the human GM affecting IDO1 expression/activity and approaches by which those can be modified to minimize this gut-inflammation-IDO1 activation, and thereby reduce its impact on gestational metabolism. Interestingly, human studies have emerged, although with unclear mechanism, that probiotics/prebiotics may minimize the development of GDM ^48^.

## Materials and Methods

### Experimental animals

Female C57BL/6J (Jackson Laboratory), CD-1 (Charles River Labs), and NIH-Swiss (Envigo) mice (age 10 weeks, n=10/strain/time-point) were individually housed in a temperature and humidity controlled specific pathogen free barrier facility with ad lib access to autoclaved food (Envigo 7912) and water. Following acclimation, pregnancy was induced using male mouse (same strain). Upon pregnancy confirmation, the male was removed to avoid contamination. Feces, cecal material, blood, liver, pancreas, and adipose tissues collected on gestation days: G0, G10, G15, G19 and post-partum days: PP3, PP20 were flash-frozen in liquid N_2_ and stored at - 80^°^C until processed. For separate set of experiments, female C57BL/6J mice (age 10-11 weeks) were given 2mg/ml IDO1 inhibitor (1-methyl-L-tryptopham, L1MT; Sigma-Aldrich) in drinking water supplemented with Splenda sweetener (2 sachets/liter) or Splenda alone as before ^49^ and female IDO1-KO mice (originally purchased from Jackson Laboratory and maintained in house) were followed during pregnancy.

To create pseudo-GF mice before FMT, C57BL/6 female mice (age 10-11 weeks) received penicillin (2000 units/ml) and streptomycin (2000μg/ml) in drinking water for 3 days. GF mice were next colonized (single 200μl gavage of inoculums prepared as before ^5, 50^) with either de-identified stool samples from T1 and T3 from 2 women (age 18-33 years, BMI 24-26.6) or with G0 and G15 fecal samples from 4 C57BL/6 mice collected earlier and monitored for 2 weeks followed by insulin tolerance test (day 14) and sac (day 15). Human sample collection protocols and mouse studies were approved by the University of Illinois at Chicago (UIC) Institutional Review Board (IRB# 2014-0325) and the IACUC of the Jesse Brown VA Medical Center and UIC, respectively and performed in accordance with the Guide for the Care and Use of Laboratory Animals.

### Metabolic assays and measurements

*In vivo* studies. Mice were fasted 16h and 6h prior to intraperitoneal glucose (2 g/kg) or insulin (1.0 U/kg) injection, for glucose and insulin tolerance tests, respectively and tests conducted as before ^50^. 16h fasted mice received 2 U/kg insulin or saline intraperitoneally euthanized 15min later. Protein lysates from harvested tissues were prepared in RIPA lysis buffer (Cell Signaling), estimated using Bradford reagent (Bio-Rad), separated on 10% SDS-PAGE (Bio-Rad) and analyzed for insulin signaling pathway using immunoblotting.

Analyte measurements: Plasma Tnfα was assayed by ELISA (Proteintech), LPS by pierce chromogenic endotoxin quant kit (Thermoscientific) and liver triglyceride by reagents from Wako Diagnostics (Richmond). Plasma samples at different pregnancy time points were assayed via LC-MS/MS using an untargeted approach as before ^51^ and via targeted approach for SCFAs ^52^ and tryptophan metabolites. For the latter, homogenized ileal tissues and plasma were deproteinized with methanol. L-kynurenine sulfate was used as internal standard. LC-MS/MS measurements were performed at the Mass Spectrometry Core facility of the UIC Research Resource Center.

#### RNA extraction and qPCR

1 μg purified extracted RNA (RNeasy Mini Kit, Qiagen) was reverse transcribed using qScript Reverse Transcriptase (Quanta Biosciences) and amplified using iTaq Universal Syber Green supermix (BioRad) with primers listed in Supplementary Table 4.

### 16S rRNA gene sequencing and data analyses

Briefly, bacterial DNA extracted from feces was used for amplification of the V4 region of the 16S rRNA gene (515 F-806R). Samples were sequenced on the Illumina MiSeq platform at the Argonne National Laboratory core sequencing facility according to EMP standard protocols (http://www.earthmicrobiome.org/emp-standard-protocols/its/). Sequence processing and analyses was conducted as before ^53^.

### Metabolomics, clustering, metabolite-microbe/health correlations and Monocle3 analyses

Feature detection and basic quantitation of raw LC-MS data was performed using the OpenMS toolkit separately for positive and negative acquisition modes ^54^. Within each strain, mean imputation was used to replace missing values and for each feature, the mean value at G0 was subtracted from all values in order to normalize each strain to a baseline. The values were then normalized to 0 mean and unit variance within each strain. K-means clustering was used across all strains using 4 clusters. Microbes predictive of the G15/G19 gestational states were identified by Meta-Signer, which is an ensemble of four machine learning models to rank microbes based on their predictive power. The top 10 microbes from the rank list were selected for further analysis.

Before differential analysis of metabolites, values were clipped to a maximum value of 3 times the metabolite’s maximum value within each strain to shrink the effect of outlier values. Metabolite values were then log-transformed and scaled to 0 mean and unit variance within each strain. Metabolites not appearing within 20% of any strain were removed. For each strain, PERMANOVA was then used to identify metabolites significantly different between GA15/19 samples and the rest with a *P*-value cut-off of .05 after Benjamini-Hochberg correction. Significantly enriched pathways for each strain were identified using the mummichog analysis in MetaboAnalyst. The candidate annotated metabolites were mapped back to the untargeted features, selecting the most significantly differential metabolite based on the original PERANOVA analysis if a candidate mapped to multiple features. Spearman correlation correlations were calculated using the top 10 microbes identified by Meta-Signer, the top 10 most significant annotated metabolites, and the set of health parameters. Benjamini-Hochberg corrected *P*-values were reported.

Monocle3 was used for trajectory analysis for each data type, where the strain specific preprocessed data were grouped into a single set. Monocle3 was run using UMAP dimension reduction and Louvain clustering.

### Statistical analyses

Microbial diversity (alpha diversity, based on Shannon and Inverse Simpson indices and beta diversity) were assessed for significance as before ^53^. The differences in alpha and beta diversity indices were then tested for significance using Kruskal-Wallis and permutational multivariate analysis of variance (PERMANOVA), respectively, with *P*-values corrected using Benjamini-Hochberg FDR correction. The analyses of composition of microbiome (ANCOM) followed by Mann-Whitney U test was used to identify differentially abundant bacterial ESVs between different groups i.e. three strains and different pregnancy stages at *P*-value cut-off of .05 and Benjamini-Hochberg FDR correction. Weighted correlation network analysis (WGCNA) package in R was used to identify clusters (modules) of significantly correlated ESVs and examine their associations with physiological variables as before ^53^.

All other data are expressed as means±SE and analyzed by Student’s two tailed unpaired *t* tests, one-way ANOVA or two-way ANOVA with post hoc tests (GraphPad Software 9.0) as applicable. *P*<.05 was considered significant.

## Abbreviations

16S rRNA: 16S ribosomal RNA
Akt: protein kinase B
ANCOM: analyses of composition of microbiome
ANOVA: analysis of variance
Cldn4: claudin 4
Cxcl1: C-X-C Motif Chemokine Ligand 1
DC: distal colon
E-cad: E-cadherin
ESV: Exact Amplicon Sequence Variant
FDR: false discovery rate
FMT: fecal microbial transfer
G: gestation day
GDM: gestational diabetes mellitus
GM: gut microbiome
IDO1: indoleamine 2,3-dioxygenase
IDO-KO: indoleamine 2,3-dioxygenase knock out
Ifnγ: interferon γ
Il: interleukin
IR: insulin resistance
K/T: kynurenine to tryptophan ratio
L1MT: L-1Methyltryptophan
LC-MS: Liquid chromatography–mass spectrometry
Lcn2: lipocalin 2
LPS: lipopolysaccharide
NMDS: non-metric multidimensional scaling
Ocln: occludin
PERMANOVA: permutational multivariate analysis of variance
PP: postpartum day
GF: pseudo germ-free
Reg: regenerating islet-derived protein
SCFA: short chain fatty acid
SE: standard error
T: trimester
Tgfβ: transforming Growth Factor Beta 1
TJ: tight junction
Tnfα: tumor necrosis factor *α*
WGCNA: Weighted correlation network analysis (WGCNA)

## Grant support

National Institutes of Health under award number R01DK104927-01A1; University of Chicago Diabetes Research and Training Center (P30DK020595); and Department of Veterans’ Affairs, Veterans Health Administration, Office of Research and Development, VA merit (grant no. 1I01BX003382-01-A1) for BTL. YD acknowledges supports from the NVIDIA Corporation with the donation of the Titan Xp GPU used for this research. RKG acknowledges NIH DK098170 (R01, NIH/NIDDK). CRM acknowledges NIH F30 DK113703. BPB, JG, PMM acknowledge R03 HD095056. BPB also acknowledges Arnold O. Beckman Postdoctoral Fellow Award and BIRCWH Training award (K12 HD101373).

## Conflict of interest

none

## Author contributions

Conceptualization: BTL, MP, GN; Data curation: MP, GN, BS, CM, DR, AS, KL, KX; Analysis: MP, GN, AS, DR, YD, GC, BW, BTL; Funding Acquisition: BTL; Investigation: MP, GN, BS, CM, KX, KL; Methodology: MP, GN, CM, AS, DR, YD, WK, RKG, BTL; Project Administration: MP, GN, BTL; Resources: BTL, JG, YD, BPB, PMM; Software: MP, GN, BW, AS, DR, YD, JG; Supervision: BTL, MP; Validation: MP, BTL; Visualization: KX, MP; Writing (original draft): MP, GN, AS, DR, GEC, BW, YD, BTL; Writing (editing and revising): MP, AS, DR, BPB, RKG, JG, BW, YD, BTL

## Data transparency statement

Data, analytic methods, and study materials will be made available to other researchers online on website associated with the lab.

---

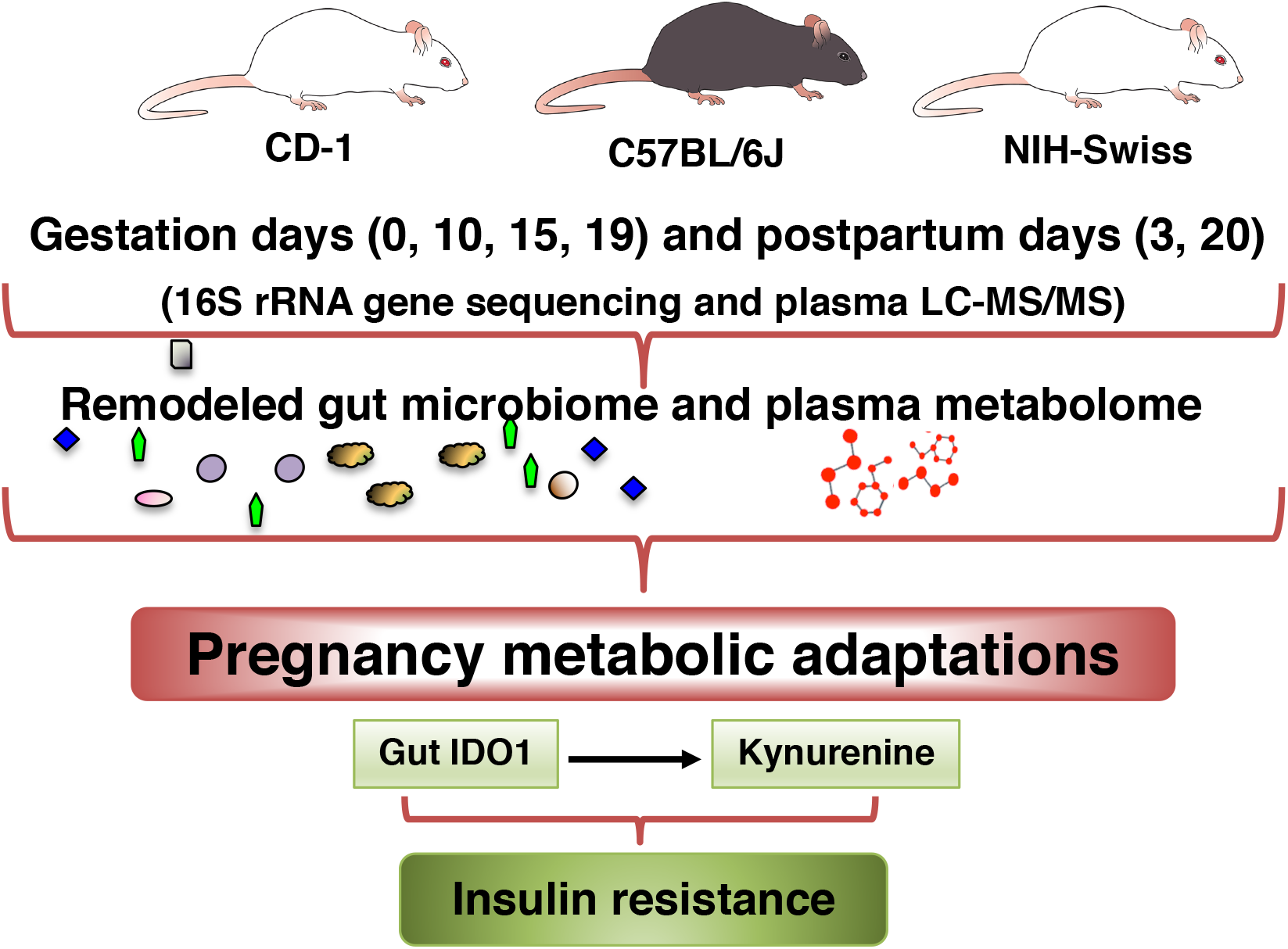

**Supplemental Figure 1.**
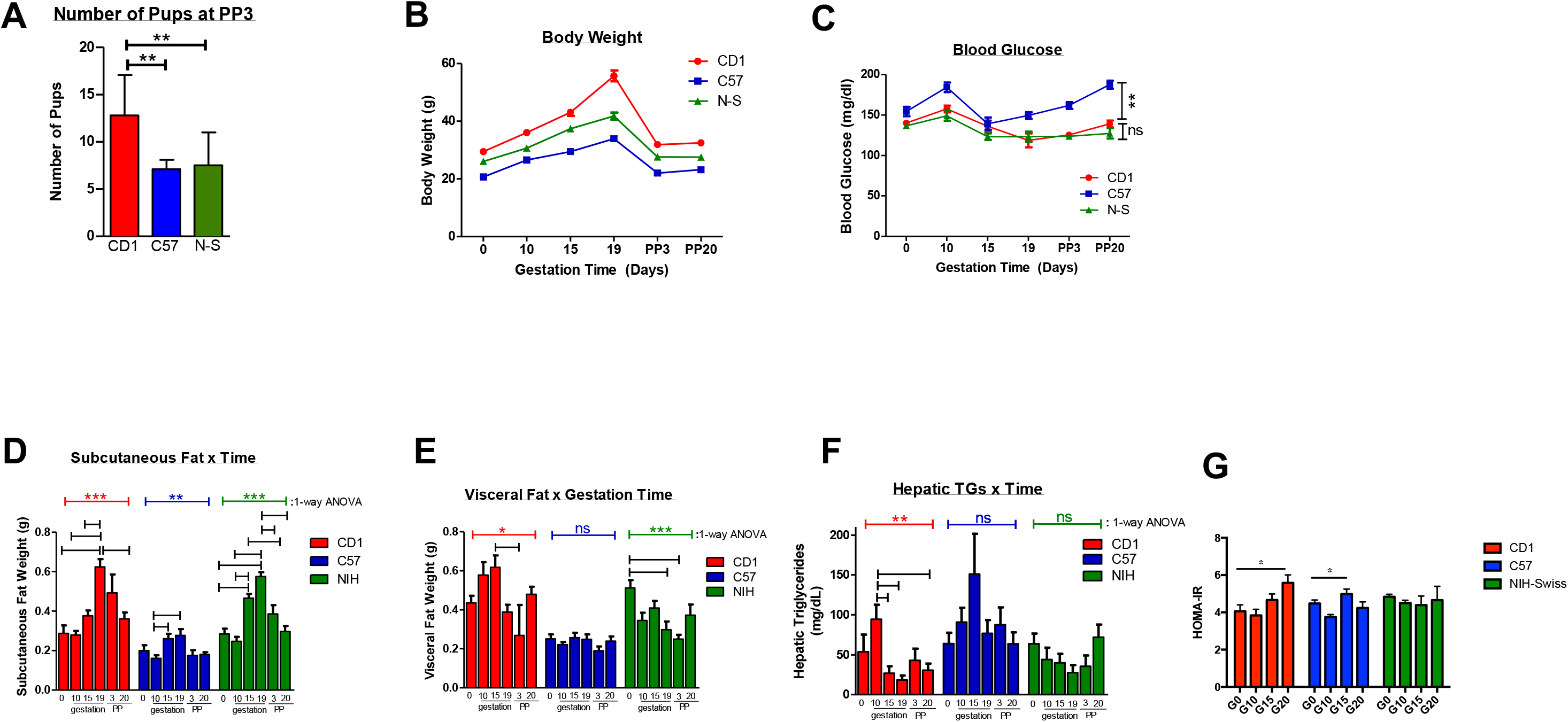

**Supplemental Figure 2.**
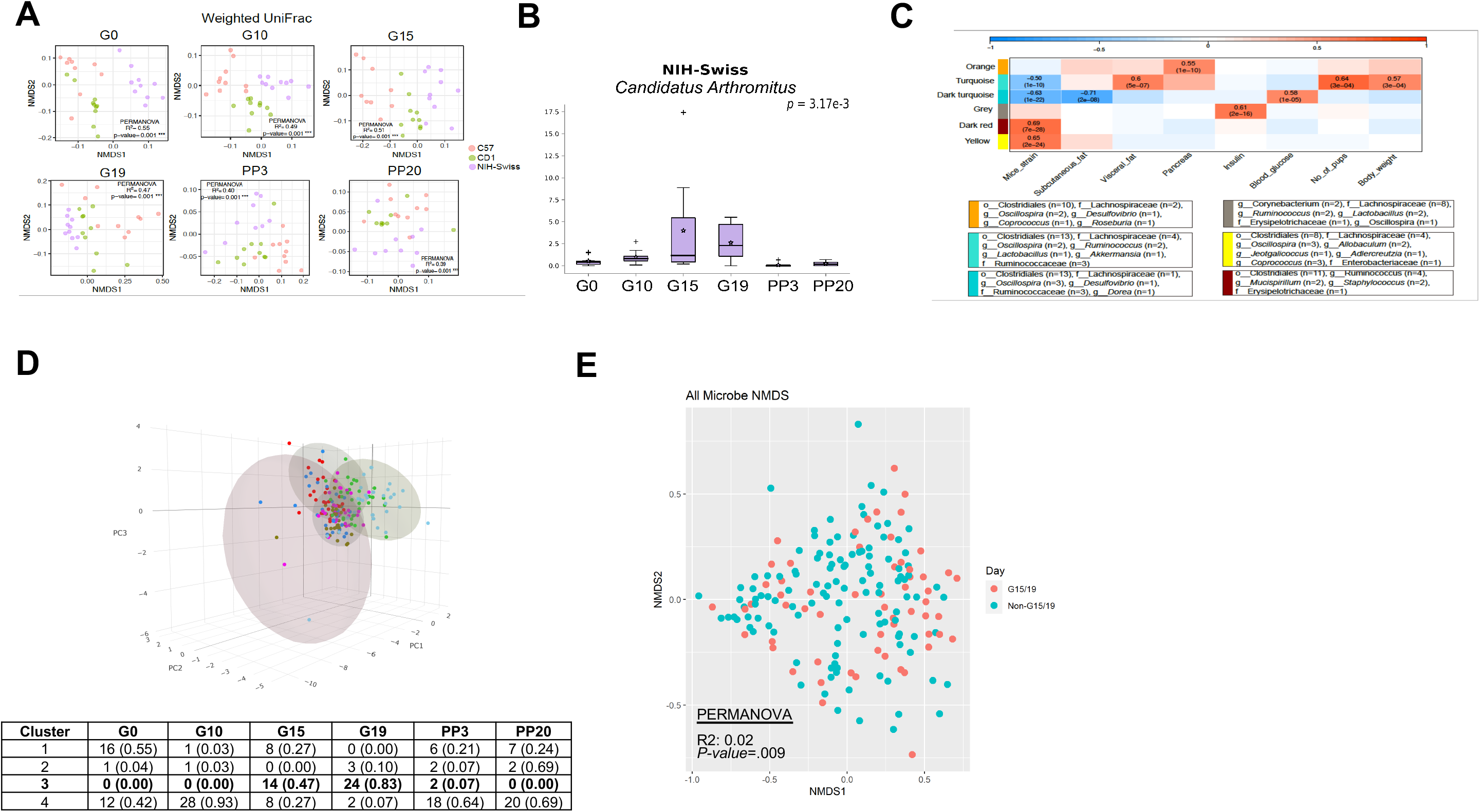

**Supplemental Figure 3.**
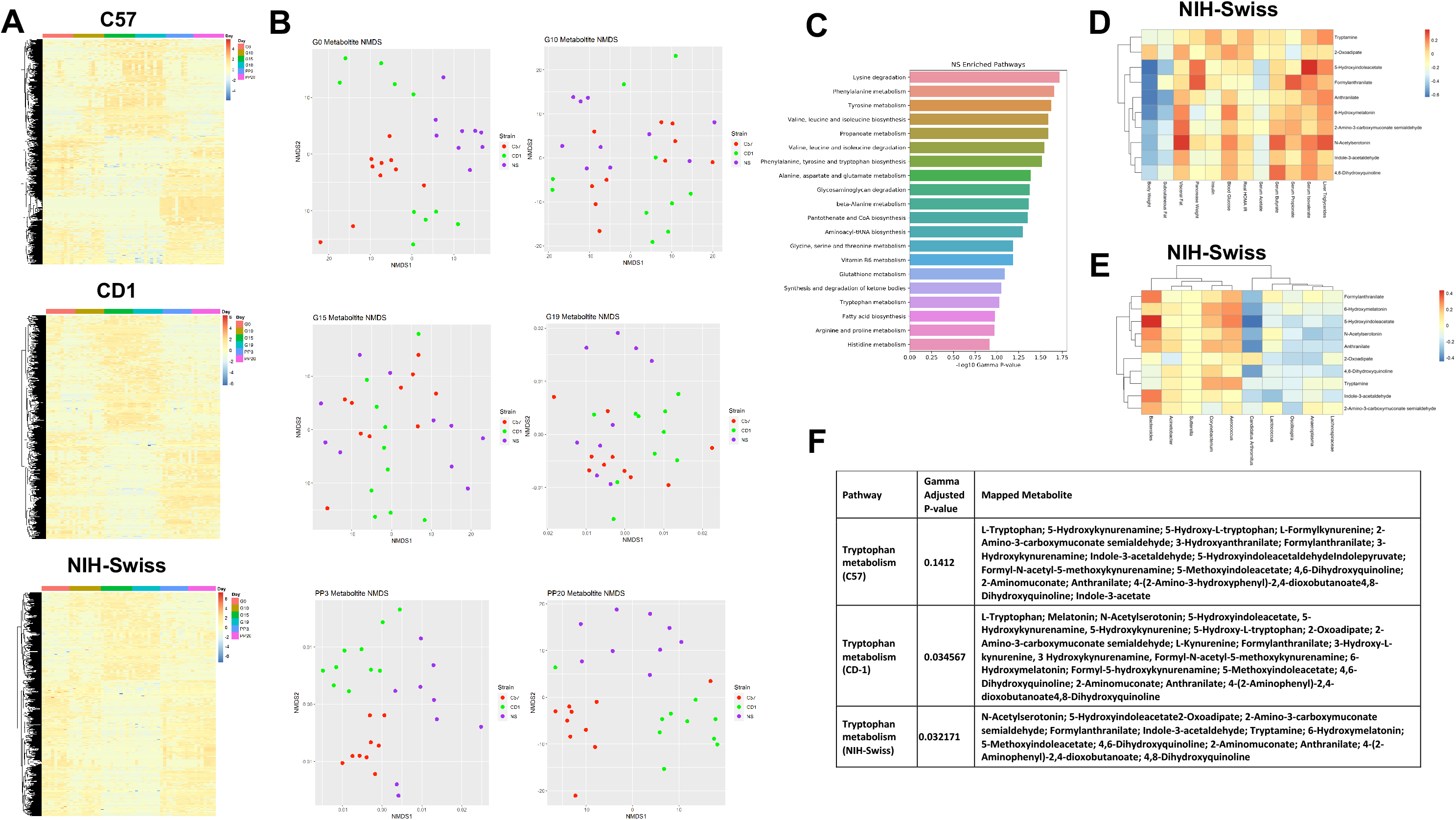

**Supplemental Figure 4.**
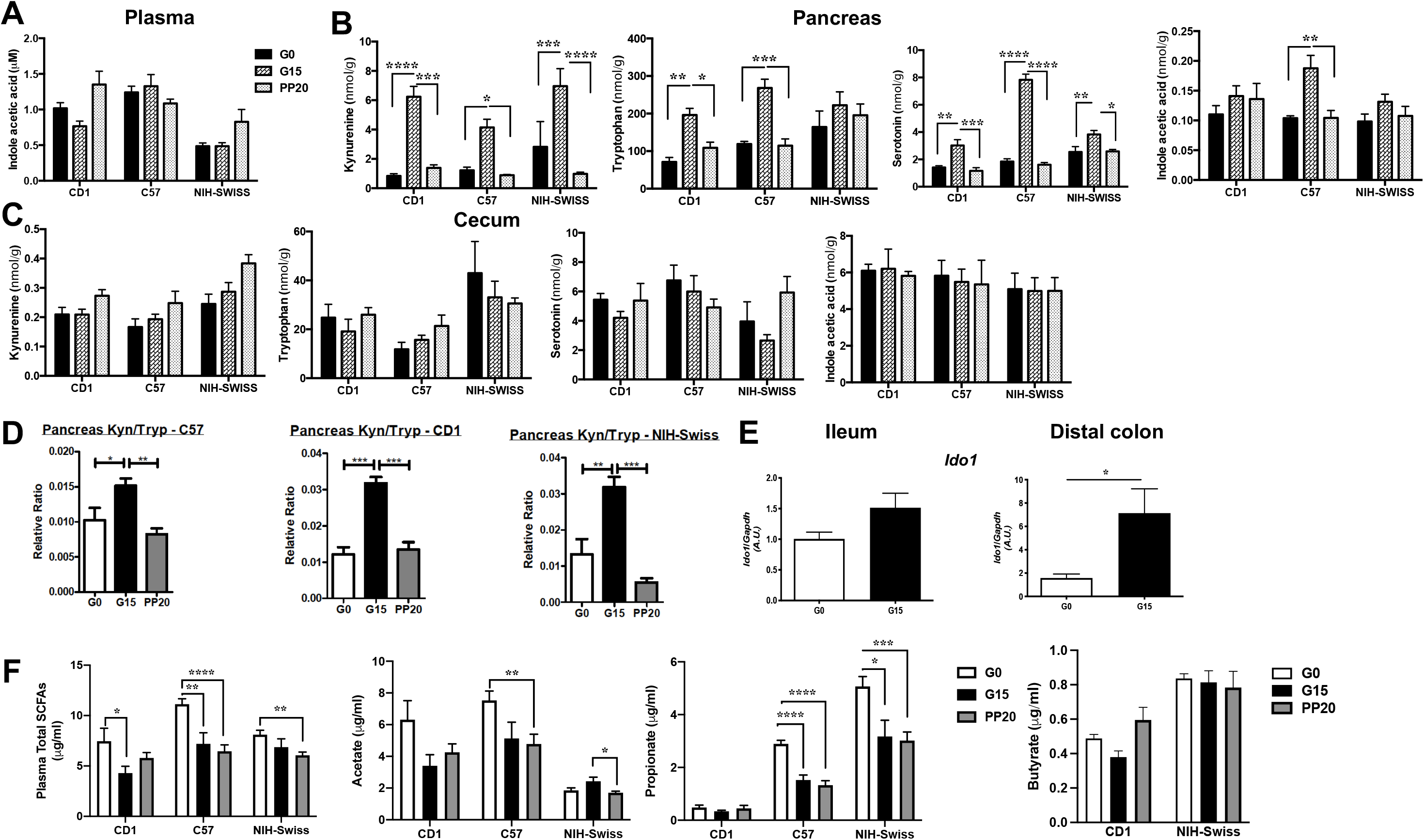

**Supplemental Figure 5.**
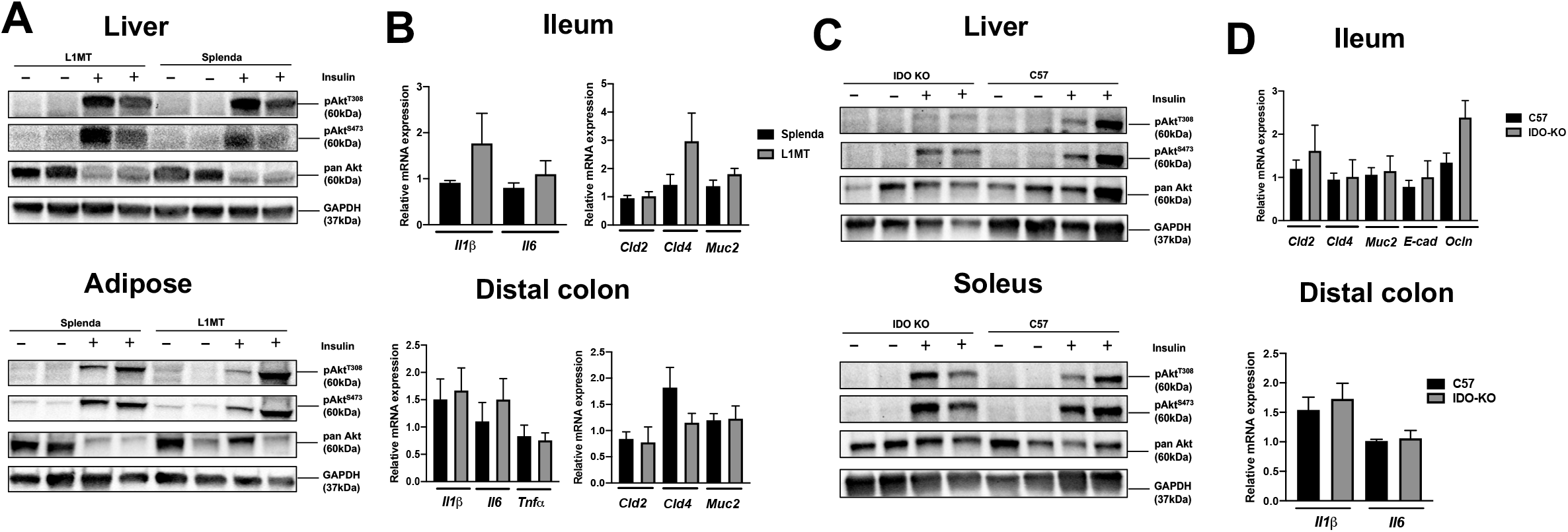

## Notes

### Competing Interest Statement

The authors have declared no competing interest.

