## Supplemental information for "Gestational insulin resistance is mediated by the gut microbiome-indoleamine 2,3-dioxygenase axis"

**Supplementary Results**

**Taxa-trait relationships**

To investigate the relationship between bacterial taxa and individual metabolic parameters (blood glucose, plasma insulin, adipose depot weight, body weight) and non-metabolic parameters (mouse strain, number of pups), correlation analyses were performed using Weighted correlation network analysis (WGCNA) in R (**Fig. S2c**). WGCNA is a network-based approach to organizing taxa into a hierarchy of modules where members in a module share similar abundance level in all samples. WGCNA was first used to identify set of highly correlated taxa (modules) and further correlation between modules and metabolic properties was estimated using eigen-networks implemented in WGCNA. The correlation network, based on the co-occurrence of the ESVs, could be divided in 5 distinct groups, communities or modules. The orange module [*Oscillospira, Desulfovibrio, Coprococcus*, *Roseburia*] positively correlated with pancreas weight (0.55, 1e-10). The dark turquoise module [*Oscillospira, Dorea*, family Lachnospiraceae, order Clostridiales] negatively correlated with mouse strain (-0.63, 1e-22) and subcutaneous fat (-0.71, 2e-08) and positively correlated with blood glucose (0.4, 0.005). The grey module [*Oscillospira, Corynebacterium, Lactobacillus, Ruminococcus*] positively correlated with plasma insulin (0.61, 2e-16). The dark red and yellow modules [*Ruminococcus, Staphylococcus, Mucispirillum, Coprococcus, Oscillospira, Allobaculum*] positively correlated with mouse strain (0.69, 7e-28 & 0.65, 2e-24, respectively). The turquoise module [*Ruminococcus, Oscillospira, Lactobacillus, Akkermansia, Proteus*] positively correlated with visceral adipose tissue weight (0.6, 6e-09), number of pups (0.57, 3e-04), and body weight (0.57, 3e-04). The bacterial genera *Oscillospira* and *Ruminococcus* were among the taxa most positively associated with the metabolic parameters and phenotypes. These data suggest a collective role for these organisms in pregnancy adaptations.

Machine learning analyses investigated relationships between microbial taxa and gestational stage across mouse strains. Clustering analysis (k-mean) using the metabolic response profiles showed that most G15/G19 mice (63.3%) shared a similar metabolic profile (**Fig. S2d**) suggesting that irrespective of the strain, later pregnancy induces a unique state of metabolic reprogramming.

Figure legends (Suppl)

**Figure S1. Physiological response of three distinct mouse strains to pregnancy.** **(a)** Number of pups; **(b)** body weight; **(c)** blood glucose; **(d-e)** weights of subcutaneous and visceral adipose tissues; **(f)** hepatic triglycerides; **(g)** HOMA-IR in C57BL/6J, CD1 and NIH-Swiss mice at different time points during gestation (G) and post-partum (PP) (n=10 for each strain per time point). Data are expressed as mean±SE and analyzed by one-way ANOVA with Tukey’s post-hoc test. **P*<.05, ***P*<.01, ****P*<.0005.

**Figure S2. Pregnancy impact on fecal bacterial diversity and select bacterial taxa. (a)** NMDS plots showing microbial beta-diversity using weighted UniFrac within each gestational stage, where a PERMANOVA analysis indicate significant separation (*P*<.001) between strains in each gestational stage; **(b)** multi-group non-parametric ANCOM test followed by Mann-Whitney U test showing differentially abundant genera at various stages of pregnancy in NIH-Swiss mice; **(c)** WGCNA analysis to assess significant associations of functional modules (clusters of Exact amplicon Sequence Variation) and a specific (physiologic) trait. Each row corresponds to a module eigengene, column to a trait. Each cell contains the corresponding correlation and *P*‐value (Benjamini-Hochberg FDR correction), n=number of ESVs from that taxa. The table is color coded by correlation according to the color legend; **(d)** PCA visualization of health parameters with *k*-means clustering using four clusters. The table shows the number of samples from each gestational stage within each cluster with the proportion of samples from a gestational stage found within a cluster is shown in parentheses; **(e)** NMDS plot using all microbes and colored by G15/19 and non-G15/G19 gestational states is shown, with a PERMANOVA analysis indicated in the plot.

**Figure S3. Pregnancy impact on metabolomic profiles.** **(a)** Heatmaps showing the differentially expressed metabolites (*P*<.05) identified by a PERMANOVA analysis between G15/19 and Non-G15/19 gestational states for C57 (top), CD1 (middle), and NIH-Swiss (bottom) strains; **(b)** NMDS plots of each gestational stage using metabolomic data; **(c)** the top 20 most significantly enriched pathways identified by mummichog using the differentially expressed metabolites between G15/19 and non-G15/19 samples in the NIH-Swiss strain (left). **(d-e)** Spearman correlation analyses between the top 10 most significant metabolites from the tryptophan metabolism pathway that were able to be mapped using mummichog candidates and both the health parameters **(d)**, the top 10 microbes identified from Meta-Signer for NIH-Swiss **(e)**; **(f)** table shows gamma adjusted *P*-values and mapped metabolites for the tryptophan metabolism pathway for each of the three strains.

**Figure S4. Modulation of tryptophan metabolites during pregnancy.** **(a)** plasma indole acetic acid (IAA); kynurenine, tryptophan, serotonin, and IAA in **(b)** pancreas, and **(c)** cecum; **(d)** Kyn/Tryptophan ratio in pancreas of C57BL/6J, CD1 and NIH-Swiss mice at G0, G15 and PP20 (n=5 per time point); **(e)** mRNA expression of *ldo1* in ileum (*left*) and distal colon (*right*) of CD1 mice; **(f)** plasma total short chain fatty acids (SCFAs), acetate and propionate levels in C57BL/6J, CD1 and NIH-Swiss mice and butyrate levels in CD1 and NIH-Swiss mice at G0, G15 and PP20 (n=10 per time point). Data are expressed as mean±SE and analyzed by one-way ANOVA with Tukey’s post-hoc test **(a-d)**, two-tailed Student’s unpaired *t* test **(e)** and two-way ANOVA (mixed effects analysis) **(f)**. **P*<.05, ***P*<.01, ****P*<.0005.

**Figure S5. Insulin signaling and gut inflammatory changes in absence of IDO1 activity and expression.** Insulin signaling (p-Akt-T308 and p-Akt-S473) in liver *(upper panel)* and subcutaneous adipose (*lower panel*) of **(a)** L1MT treated and control mice, and **(c)** in liver *(upper panel)* and soleus (*lower panel*) of IDO-KO and control mice, at G15 (n=4-5); mRNA expression of proinflammatory mediators and gut proteins ileum *(upper panel)* and distal colon (*lower panel*) of **(b)** L1MT treated and **(d)** of IDO KO and respective control mice at G15 (n=5-7). Data are expressed as mean±SE and analyzed by Two-tailed Student’s unpaired *t* test.

**Table S1**

Differentially enriched pathways based on the mummichog analyses on the differential metabolites for C57BL/6J mice.

**Table S2**

Differentially enriched pathways based on the mummichog analyses on the differential metabolites for CD1 mice.

**Table S3**

Differentially enriched pathways based on the mummichog analyses on the differential metabolites for NIH-Swiss mice.

**Methods**

Immunoblotting for insulin signaling pathway. 30 μg of denatured proteins were separated on 10% SDS-PAGE gels (Bio-Rad Laboratories) and transferred to PVDF membranes, probed using pAkt Ser473 (CST, D9E), pAkt Thr 308 (CST, D25E6), pan Akt (CST, C67E7), GAPDH (CST, 14C10), all used at dilution of 1:1000.

**Supplementary Table 4**. Primer sequences used for qPCR

| **Gene** | **Forward Primer 5’-3’** | **Reverse Primer 5’-3’** |
| --- | --- | --- |
| *Ido1* | CAAAGCAATCCCCACTGTATCC | ACAAAGTCACGCATCCTCTTAAA |
| Inflammatory markers | | |
| *Ifng* | ATGAACGCTACACACTGCATC | CCATCCTTTTGCCAGTTCCTC |
| *Cxcl1* | AAAGATGCTAAAAGGTGTCCCCA | AATTGTATAGTGTTGTCAGAAGCCA |
| *Il1b* | GCAACTGTTCCTGAACTCAACT | ATCTTTTGGGGTCCGTCAACT |
| *Il6* | TAGTCCTTCCTACCCCAATTTCC | TTGGTCCTTAGCCACTCCTTC |
| *Il12* | AGACCCTGCCCATTGAACTG | GAAGCTGGTGCTGTAGTTCTCATATTT |
| *Tnfa* | GGTGCCTATGTCTCAGCCTCTT | GCCATAGAACTGATGAGAGGGAG |
| Anti-inflammatory markers | | |
| *Il22* | ATGAGTTTTTCCCTTATGGGGAC | GCTGGAAGTTGGACACCTCAA |
| *Il10* | CGCAGCTCTAGGAGCATGTG | GCTCTTACTGACTGGCATGAG |
| *Tgfb* | CTGAGTGGCTGTCTTTTGACG | CGTGGAGTTTGTTATCTTTGCTG |
| Gut barrier markers | | |
| *Ecad* | CAGCCTTCTTTTCGGAAGACT | GGTAGACAGCTCCCTATGACTG |
| *Ocln* | CCTCCAATGGCAAAGTGAAT | CTCCCCACCTGTCGTGTAGT |
| *Muc2* | GTGTGGGACCTGACAATGTG | ACAACGAGGTAGGTGCCATC |
| *Cld2* | TTAGCCCTGACCGAGAAAGA | AAAGGACCTCTCTGGTGCTG |
| *Cld4* | AGCAAACGTCCACTGTCCTT | AATCCACCTCCACCCTTCTT |
| Antimicrobial markers | | |
| *Lcn2* | CCTCCATCCTGGTCAGGGAC | TAGTCCGTGGTGGCCACTTG |
| *Reg3b* | ATGCTGCTCTCCTGCCTGATG | CTAATGCGTGCGGAGGGTATATTC |
| *Reg3g* | GTTGCCAAGAAAGATGCCCC | GGCAGGCCATATCTGCATCA |
| housekeeping | | |
| *Gapdh* | agcttgtcatcaacgggaag | tttgatgttagtggggtctcg |
